# Single-cell transcription profiles in Bloom syndrome patients link *BLM* deficiency with altered condensin complex expression signatures

**DOI:** 10.1101/2021.10.01.462717

**Authors:** Ipek Ilgin Gönenc, Alexander Wolff, Julia Schmidt, Arne Zibat, Christian Müller, Lukas Cyganek, Loukas Argyriou, Markus Räschle, Gökhan Yigit, Bernd Wollnik

## Abstract

Bloom syndrome (BS) is an autosomal recessive disease clinically characterized by primary microcephaly, growth deficiency, immunodeficiency, and predisposition to cancer. It is mainly caused by biallelic loss-of-function mutations in the *BLM* gene, which encodes the BLM helicase, acting in DNA replication and repair processes. Here, we describe the gene expression profiles of three BS fibroblast cell lines harboring causative, biallelic truncating mutations obtained by single-cell (sc) transcriptome analysis. We compared the scRNA transcription profiles from three BS patient cell lines to two age-matched wild-type controls and observed specific deregulation of gene sets related to the molecular processes characteristically affected in BS, such as mitosis, chromosome segregation, cell cycle regulation, and genomic instability. We also found specific upregulation of genes of the Fanconi anemia pathway, in particular *FANCM, FANCD2*, and *FANCI*, which encode known interaction partners of BLM. The significant deregulation of genes associated with inherited forms of primary microcephaly observed in our study might explain in part the molecular pathogenesis of microcephaly in BS, one of the main clinical characteristics in patients. Finally, our data provide first evidence of a novel link between BLM dysfunction and transcriptional changes in condensin complex I and II genes. Overall, our study provides novel insights into gene expression profiles in BS on a single-cell level, linking specific genes and pathways to BLM dysfunction.

## Introduction

Bloom syndrome (BS, MIM: 210900) is a rare genetic disease that was first described in 1954 in children with growth retardation and sunlight sensitivity (1). BS is defined as an autosomal recessive disease with clinical characteristics of primary microcephaly, pre- and postnatal growth deficiency, short stature, immunodeficiency, and a predisposition to cancer with an early age of onset (2–4). A narrow face with a sunlight-sensitive erythematous rash on the cheeks and café-au-lait spots are the differential characteristics of BS (1,5). Mainly, biallelic loss-of-function mutations in *BLM* including nonsense and missense variants as well as frameshifting indels cause the BS phenotype (4). The *BLM* gene encodes the BLM RecQ-like helicase, an enzyme that unwinds double-stranded DNA during DNA replication and repair processes (6). Furthermore, the BLM helicase forms a part of a multiprotein complex called BTR complex, which also includes topoisomerase III alpha (TOP3A) and RecQ-mediated genome instability protein 1 (RMI1) and 2 (RMI2) (7–9). Recently, homozygous frameshift mutations in *TOP3A* and *RMI1* as well as loss of the *RMI2* gene as a result of a large homozygous deletion have been linked to a BS phenotype (10,11).

The BTR complex plays important roles in the maintenance of genome stability, dissolving DNA intermediates such as G-quartets, D-loops, and Holliday junctions (12). The BLM helicase moves the branches of double Holliday junctions towards each other and the topoisomerase III alpha unhooks the hemicatenated structure, resulting in non-crossover products (13,14). For now, the BTR complex is the only known protein complex that is able to dissolve the double Holliday junctions in such a manner that no exchange between sister chromatids occurs during the homology-directed repair process (15,16). Further actions of the BTR complex such as localization to stalled replication forks and DNA end resection during double-stranded break (DSB) repair emphasize its importance for the protection of the genetic material (17). The cellular phenotypes of BS related to genomic instability include, e.g., mitotic aberrations such as chromatid breaks, anaphase bridges, and high levels of somatic mutations (11,16,18). Gene expression levels are also affected in BS, but studies investigating the transcriptional profile alterations in BLM deficiency are rare. One recent study examined the expression profiles of BS fibroblasts by means of RNA microarray and found out that the expression levels of the genes with G4 motifs were deregulated in BS cell lines (19). Another recent study investigated the expression levels in BS by microarray analysis and emphasized the regulation of inflammatory interferon-stimulated genes in BS cells (20). Montenegro et al. performed RNA-seq for two BS patients and reported abnormal expression levels of immune response and apoptosis-associated genes (21). Here, we report a comprehensive study of transcription profiles of BS patient fibroblast cell lines obtained by single-cell (sc) transcriptome sequencing. We define specific deregulated gene expression levels as well as specific pathway alterations in BS cells and highlight the power of single-cell transcriptomics for elucidating the molecular pathogenesis in BS.

## Results

Two age-matched wild-type (WT) control fibroblast cell lines (GM08398 and GM00409) and three BS patient fibroblast cell lines (GM02520, GM02932, and GM02548; purchased from Coriell Institute for Medical Research) were used for scRNA profiling. The causative mutations were confirmed using Sanger sequencing on DNA isolated from each cell line (Supplementary Fig. S1). For single-cell transcriptome sequencing (scRNAseq), single cells were dispensed to a 5184-well microchip using ICELL8 system (Takara Bio) (22). We prepared libraries using the Nextera XT DNA library preparation kit (Illumina) and sequenced on an Illumina HiSeq4000.

### Quality control of single-cell transcriptome data

To determine the differences in transcriptional profiles between the BS patient cells and WT cells, we performed scRNAseq on three BS patient fibroblast cell lines and two WT fibroblast cell lines (Supplementary Table 1). Quality control (QC) of the raw data was carried out by the CogentAP pipeline and a total of 517 out of 653 cells passed the initial QC filter. 70.4% of the total reads mapped to exonic reads (Fig. 1A). The mean of the read counts per cell varied between 190,586 and 244,819 reads (Fig. 1B) and the number of expressed genes per cell was approximately 7,000 for each cell line (Fig. 1C). In addition, the level of ribosomal RNA contamination in the raw data was low (almost zero), as well as the level of mitochondrial RNA percentage, which was below our set quality target of 10%. (Supplementary Figure S2). Next, the dimension reduction of the data was done by using UMAP (Uniform Manifold Approximation and Projection) (23,24) (Fig. 1D). The UMAP analysis displayed that cells belonging to the same cell line were clustering at close positions in the space and had similar expression profiles, while cells from different cell lines were distinguishable in terms of their localization in the UMAP and their expression profiles (Fig. 1D). Together, these results showed that the quality of the obtained scRNAseq data was robust and suitable for further analysis of transcriptional profiles of BS and WT fibroblasts on a single-cell level.

**Figure 1.**
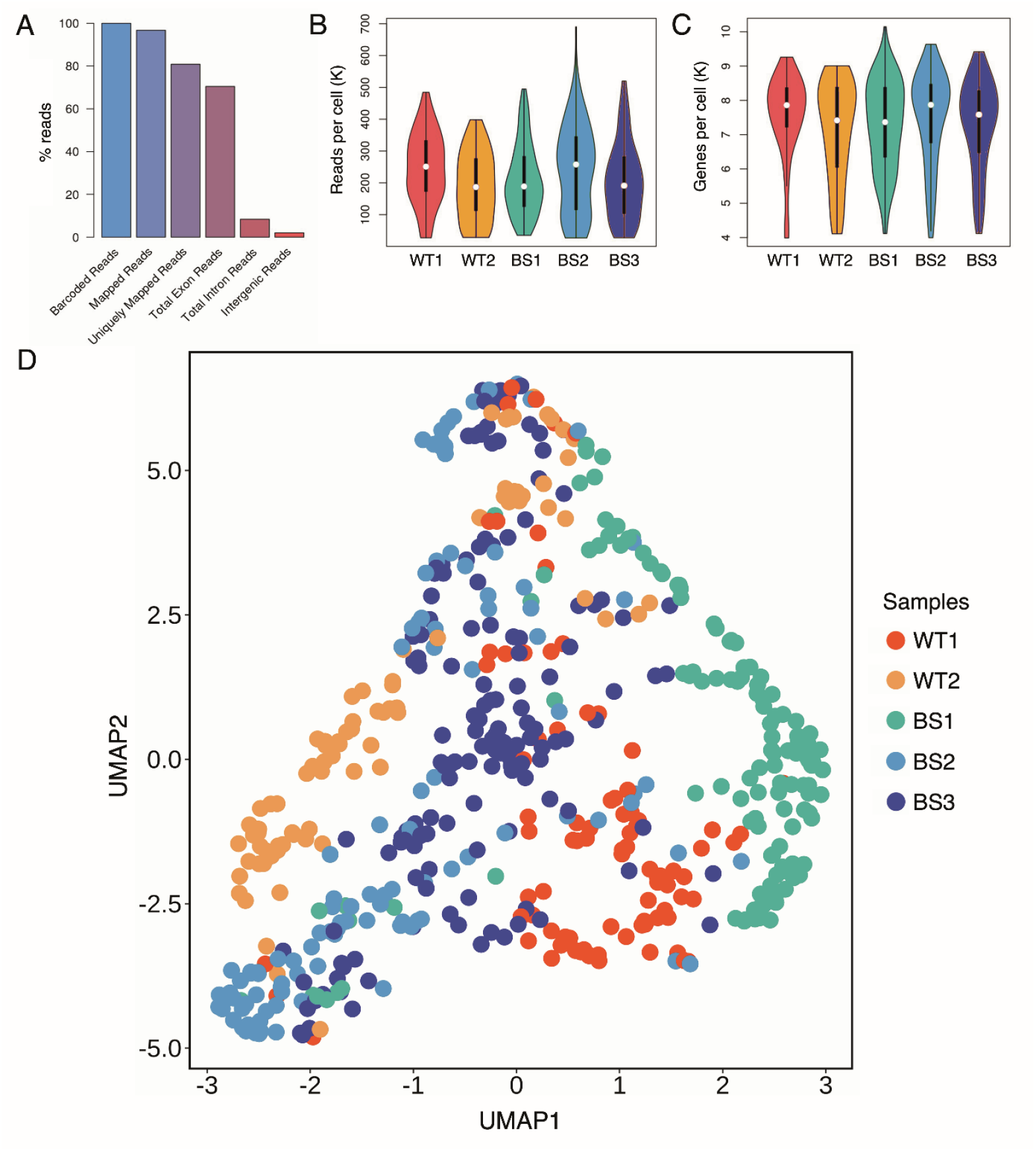
Quality control elements obtained from the CogentAP pipeline (Cogent NGS Analysis Pipeline, Takara Bio) reveal a robust single-cell transcriptome sequencing technique. (A) More than 70% of all reads mapped to exonic reads. (B) After removing cells with less than 10,000 total reads or less than 300 genes, the number of reads per cell was around 250K for all samples including two wild type (WT1, WT2) and three BS patient fibroblast cell lines (BS1, BS2, BS3). (C) After removing genes with less than 100 total reads or expressed in fewer than 3 cells, the overall gene number per cell was between 7,000 and 8,000 genes for all samples. (D) The UMAP analysis of raw data presents the different gene expression profiles of each cell line originating from healthy and patient cell lines. Clustering between cells from different cell lines can be observed on the UMAP due to differential expression profiles of single cells. The plot was generated using ggplot2 package for R-programming language (72).

### scRNAseq data confirm molecular characteristics of BS

BS patients were either homozygous or compound heterozygous for loss-of-function mutations in the *BLM* gene (Supplementary Figure S1) and the cell lines of WT controls were age-matched (Supplementary Table 1). We compared scRNAseq data from fibroblasts of three BS patient fibroblast in comparison to two WT controls and analyzed the results via over-representation analysis (ORA) for various gene sets. ORA for gene ontology (GO) terms revealed highly significant terms related to molecular signatures of BS such as mitotic defects and chromosomal aberrations (25). Strikingly, the detected deregulated terms amongst the top 20 most significant GO terms were mainly related to BS pathogenesis (Fig. 2A). The GO term with the highest significance level was “chromosome segregation” with more than 100 deregulated genes in the group, followed by “mitotic nuclear division” and “mitotic sister chromatid segregation” classes as the second and third most significant GO terms, respectively. Overall, ORA for GO terms from single cells revealed highly deregulated pathways in the BS cells in line with the molecular characteristics of the disease.

**Figure 2.**
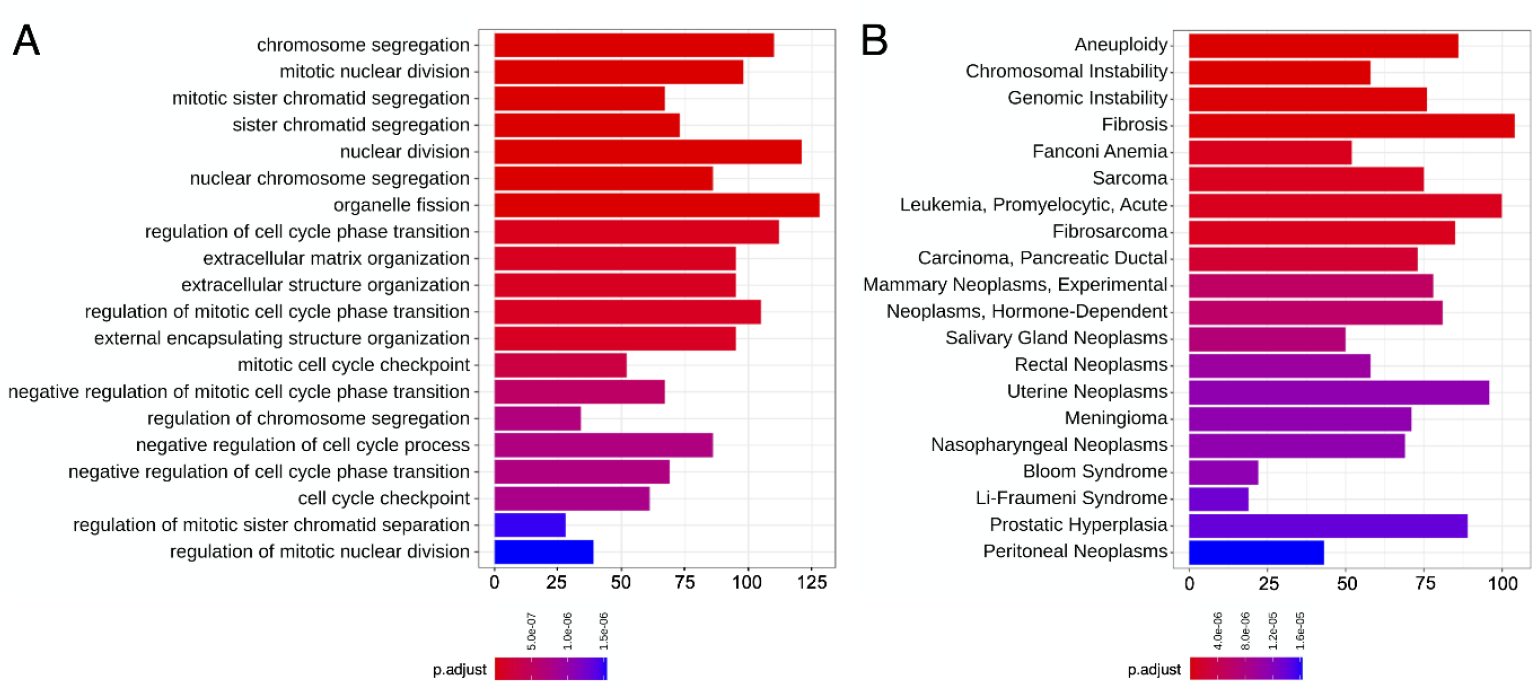
Barplots of over-representation analysis (ORA) show mainly BS-related terms in the top 20 denominations when the gene expressions of the BS patient fibroblasts and the wild-type fibroblasts were compared. (A) Top 20 differentially expressed gene ontology (GO) terms were obtained via ORA from the R-package clusterProfiler (71). (B) The top 20 Medical Subject Headings (MeSH) (26) obtained by over-representation analysis as in (A). Most of the GO terms and all of the top 20 MeSH nominations were linked to BLM dysfunction and BS pathogenesis.The x-axis shows the number of significantly deregulated genes on each denomination. The adjusted p-value calculation was done by using the Benjamini-Hochberg procedure for each p-value (70).

Next, we performed ORA using the Medical Subject Headings (MeSH) (26) to determine the association of the deregulated genes with disease-related terms. The top 20 headings with the lowest adjusted p-value contained terms relating to BS pathogenesis, such as genomic instability and various cancer-associated terms (Fig. 2B). In addition, we observed Bloom syndrome as a detected MeSH, which can be interpreted as an additional confirmation of the respective cell lines. The first three most significant terms of this analysis were “aneuploidy”, “chromosomal instability”, and “genomic instability”, each with more than 50 deregulated genes. Considering the cellular function of the BLM helicase in DNA replication and repair processes, deregulated gene expression levels of the three highest--ranking MeSH terms (aneuploidy, chromosomal, and genomic instability) were expected outcomes obtained by the single-cell transcriptome data of BS fibroblasts.

Moreover, we observed that all of the top 20 terms of the MeSH-based analysis were cancer-related terms and included cancer predisposition syndromes such as Fanconi anemia and Li-Fraumeni syndrome (27,28). One of the major and well-defined clinical characteristics of BS is the predisposition to different cancer types with a mean age of onset of 24 years (3). Our results obtained from MeSH analysis of single-cell expression profiles were in line with the clinical synopsis of BS, providing a molecular link between gene expression levels and clinical cancer predisposition characteristics of patients with BS.

### Genes of the Fanconi anemia pathway are enriched in BS

We scrutinized the deregulated pathways in relation to known functions of the BLM helicase. BLM helicase is known to act on DNA intermediates that occur during DNA replication and repair processes such as the homologous recombination pathway (15). The function of BLM in these pathways has been widely studied as a part of the BTR complex (16,29). Interestingly, none of the expression levels of members of the BTR complex were significantly deregulated in our scRNAseq data, arguing that dysregulation of *BLM* expression does not significantly affect the expression levels of the other members of the BTR complex, namely, *TOP3A, RMI1*, and *RMI2* (Supplementary Table 2).

We then investigated the expression levels of genes of the Fanconi anemia (FA) pathway, which was already detected by ORA for MeSH as a highly deregulated pathway (Fig. 2B). Mutations in the various genes of the FA pathway are linked to the autosomal recessively inherited FA disorder (30), and, interestingly, FA shows overlapping clinical features with BS such as primary microcephaly, growth deficiency, and cancer predisposition (31). On the molecular level, FA and FA-associated proteins take part in cellular processes such as DNA replication, DSB repair, and replication fork maintenance (32). It is well established that various FANC proteins directly interact with BLM during these processes (33–35). One of the important interaction partners of BLM is FANCM at stalled replication forks (34) and another important interaction of BLM was observed with BRCA1, e.g. at the telomeres of chromosomes (36).Therefore, it should be noted that both of these genes had higher expression levels in BS cells compared to WT controls (Fig. 4).

We continued our analysis by comparing the expression levels of each gene of the FA group and we observed several significantly deregulated genes of the FA pathway (Fig. 3). Interestingly, almost all of the significantly deregulated genes had higher expression levels in the BS fibroblasts compared to control fibroblasts (Supplementary Table 3). Additionally, *FANCD2* showed the largest difference in expression levels among the FA-related genes, followed by *FANCI* expression level (Supplementary Table 3).

**Figure 3.**
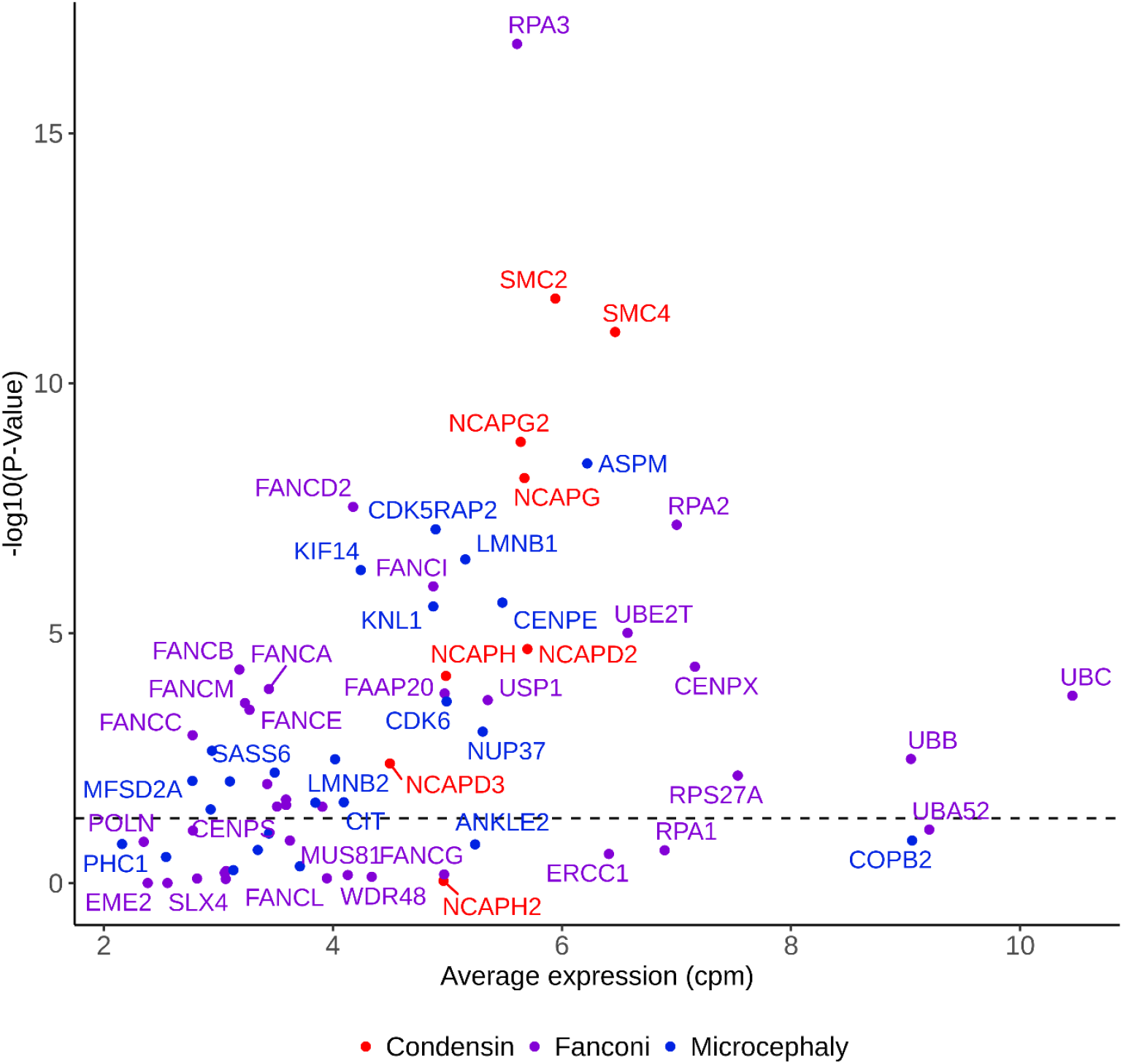
Scatterplot of gene expression shows that most of the genes in the condensin complex I/II, Fanconi anemia pathway, and the genes known to cause microcephaly (41) are significantly deregulated in BS patient cell lines in comparison to wild-type fibroblast cell lines. The color codes are as follows: condensin complex I/II genes, red; Fanconi anemia pathway genes, purple; microcephaly-related genes, blue. Label of the x-axis is the average counts per million (cpm) of each individual gene based on the raw counts. Complete gene list and the corresponding values are given in Supplementary Table 3.

### Condensin complex I and II genes are overexpressed in BS cells

Condensin complexes are ring-like multiprotein complexes that organize the DNA and chromatin in loops to form the cylinder-shaped mitotic chromosomes (37,38). Thereby, condensin complexes are important for, e.g., maintaining chromosomal stability, correct segregation of chromatids, and transcriptional control (39). Condensin complexes I and II are composed of the SMC proteins (SMC2 and SMC4) and the non-SMC proteins. Our single-cell transcriptome data analysis surprisingly revealed a highly significant upregulation of the genes coding for either condensin I or condensin II complexes, namely, *SMC2, SMC4, NCAPG, NCAPG2, NCAPH, NCAPD2*, and *NCAPD3* (Fig. 3, red color). *NCAPH2*, which encodes a condensin-II-complex-associated protein, was not significantly altered in our data, however, we interpret this result as a putative gene dropout due to the single-cell analysis (40). The gene expression levels of most of the members of condensin I and II complexes were approximately two times higher in BS cells than in control cells and the changes were highly significant (Supplementary Table 3). Based on our transcriptome data obtained from single cells, we hypothesize a link between BS and expression of condensin complex genes. In all BS samples, we observed higher expression levels of *SMC2* and *SMC4* (Fig. 4, bottom row) as well as of other members of the condensin complexes I and II. As a further experimental proof, we observed increased SMC2 and SMC4 protein levels in Western blots (Supplementary Figure S3). Overall, we suggest a molecular connection between condensation of the chromosomes and BLM function, which can be investigated in detail in future studies.

**Figure 4.**
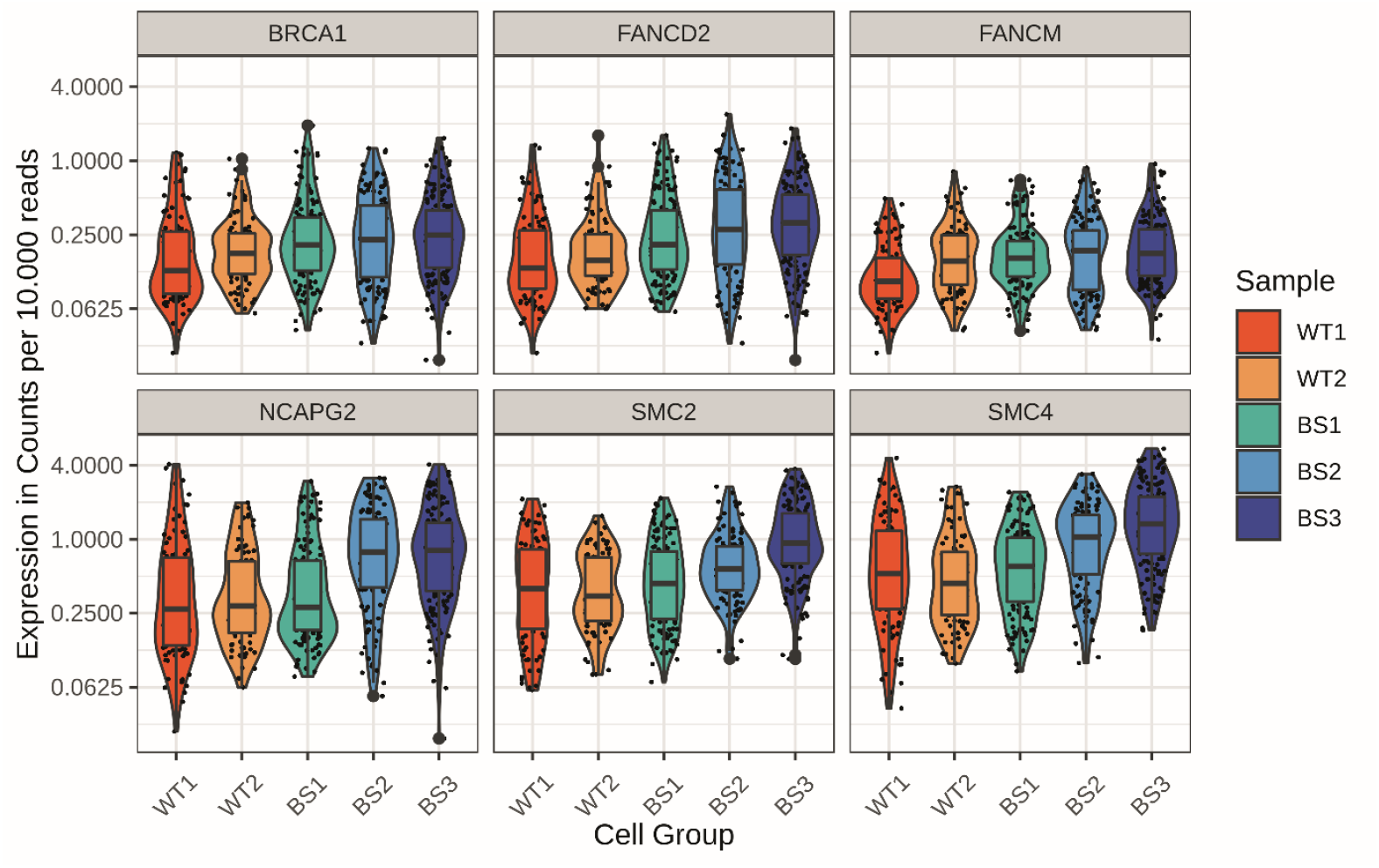
Gene expression levels (counts per 10,000) of each individual sample are shown for different genes on a single-cell level. From left to right; on the upper row *BRCA1, FANCD2*, and *FANCM*; and on the bottom row *NCAPG2, SMC2*, and *SMC4* genes are shown. The expression level differences of shown genes were significant after multiple test adjustments and the adjusted p-values are given in Supplementary Table 2. For the corresponding genes, the median expression levels were higher in BS than in WT. Visualization was done via R-package ggplot2 (72).

### Genes associated with primary microcephaly are upregulated in BS

One of the main clinical manifestations in BS is primary microcephaly. We therefore analyzed in detail the expression levels of genes which have been previously associated with autosomal recessive forms of primary microcephaly (MCPH). As shown in Fig. 3, we found altered expression levels in a significant number of MCPH-associated genes, including *CDK5RAP2, CENPE, SASS6*, and *ASPM*. Among the 27 microcephaly-associated genes (41), significant differences in expression levels were observed in 12 genes (Fig. 3, blue color, Supplementary Table 3). In addition, also mutations in genes encoding condensin I and II complex members are associated with primary microcephaly, e.g., *NCAPD2, NCAPH, NCAPG2*, or *NCAPD3*, which showed altered expression levels in our data as well (42,43).

## Discussion

Technical advances in single-cell transcriptomics offer the opportunity to determine in more detail changes in transcriptional signatures in human disease and, thereby, to gain deeper insights into the underlying disease-causing mechanisms. In this study, we systematically analyzed transcription signatures and molecular pathways in single-cell RNA sequencing in primary dermal fibroblasts from three patients with Bloom syndrome. The performance of the ICELL8 (22) platform used in our analysis allowed the investigation of a high number of single cells passing our high-quality standards. Considering all the objectives, the Limma + Voom method used for analyzing the scRNAseq data was the most effective analysis tool among other bulk and single-cell RNA sequencing analysis models (44,45). We based our analysis on three patient cell lines compared to two age-matched WT controls to receive statistically significant results. We defined specifically deregulated gene expression levels as well as significantly altered pathways in BS cells, showing the power of single-cell transcriptomics for elucidating the molecular pathogenesis in BS. Associated pathways include previously described processes relating to, e.g., mitosis, chromosome segregation, and cancer.

As described, we confirmed the deregulation of FA pathway genes in BS. This represents not only a molecular link between BS and FA, but also explains the clinical overlap between both syndromes. It is well established that several FANC proteins can directly interact with BLM during different processes (33–35), among them FANCM and FANCD2 (34,46), both of which showed altered expression in our analysis. *FANCD2* has a well-described function during DNA repair as well as an important role in replication fork management and stabilization, which implies its independent roles from the rest of the FA-related genes and proteins. Also, FANCD2 has been shown to localize to ultrafine DNA bridges, which occur frequently in BS patient cells due to the high rate of mutations and resulting replication stress (47,48). Regarding the genomic instability present in BS cells, we suggest that the enrichment of the FA pathway might be due to increased replication fork stalling and destabilization resulting from the high rate of somatic random mutations in the genome based on the affected DNA repair mechanisms (3,46,49).

Autosomal recessive primary microcephaly (MCPH) is a rare disorder mainly characterized by severe microcephaly at birth and mental retardation of variable degree in the absence of any additional significant neurological findings, malformations, or growth anomalies (50). MCPH is a very heterogeneous disorder and numerous genes have been identified which, on a cellular level, play an important role during cell division processes, regulation of the cell cycle, and in DNA damage response (51–53). Since BS patients also present with a syndromic form of primary microcephaly, we determined the expression profiles of 27 genes that were previously associated with microcephaly. A substantial number of 12 of these genes were deregulated, most significantly *ASPM*, a well-characterized MCPH gene, and *CDK5RAP2*, mutations in which have been reported in patients with primary microcephaly and/or Seckel syndrome (54–56). *CDK5RAP2* encodes the centrosomal protein CDK5RAP2, which was shown to have an important role in centriole cohesion, microtubule dynamics, and spindle orientation (57,58). Previously, we showed that loss of functional CDK5RAP2 in human fibroblasts leads to severe mitotic defects including fragmented centrosomes, unbalanced amounts of centrin and pericentrin, and multiple or misplaced centrosomes (56). The analysis of brain organoids from a patient with Seckel syndrome and homozygous *CDK5RAP2* mutation determined defects in neurogenesis caused by an altered balance of symmetric and asymmetric cell division of neuronal precursor cells (59,60). Overall, our scRNAseq data analysis strongly points towards a common or at least overlapping molecular pathogenesis of microcephaly in BS and MCPH.

Most interestingly, we observed a robust link between BLM deficiency and transcriptional upregulation of condensin complex I and II genes. This link has not yet been established. Condensins are multi-subunit SMC-containing protein complexes found in all eukaryotes (61). They are built upon specific pairs of long coiled-coil subunits that heterodimerize by the central ‘hinge’ domain situated at one end of the coil. Additional subunits are recruited to the condensin complex and, depending on the proteins associated to the complex, condensin I and II complexes are defined (62). We observed a significant upregulation of *SMC2* and *SMC4* as well as other complex subunits (*NCAPG2, NCAPG, NCAPH, NCAPD2*, and *NCAPD3*). Condensin complexes are essential for the structural organization of mitotic and meiotic chromosomes. Studies have shown that chromosome assembly is impaired when condensin subunits are dysfunctional or depleted. Cells attempt to undergo anaphase but chromosomes frequently fail to segregate, leading to lagging chromosomes and the formation of chromatin bridges (42,63). In humans, autosomal recessive mutations in genes encoding condensin complex proteins have been associated with microcephalic syndromes (42). It is of further interest that recently described recessive loss-of-function mutations in *NCAPG2* cause a neurodevelopmental disorder with microcephaly, and analysis of patient fibroblasts found not only abnormal chromosome condensation, but also augmented anaphase chromatin-bridge formation, similar to the anaphase-bridge formation observed in patients with BS and BLM dysfunction (43). On the other hand, BS cells lacking resolvases show strong condensation defects, especially at the common fragile sites resulting from unresolved recombination products which then lead to ultrafine anaphase bridges (18,64,65). Our results will have to be confirmed by future functional studies aiming to elucidate the functional interaction of BLM and condensin complexes and it will be also of interest to investigate the expression and function of *BLM* in patients with different forms of condensinopathies.

In summary, we have defined gene expression profiles and pathway signatures in patients with Bloom syndrome caused by BLM deficiency using single-cell transcriptome analysis. We observed deregulation of specific gene sets related to the molecular characteristics of BS, such as mitosis, chromosome segregation, cell cycle regulation, and genomic instability. In addition, we found specific and already described upregulation of BLM-interacting proteins, e.g., related to the Fanconi anemia pathway. Significant deregulation of genes associated with inherited forms of primary microcephaly might explain - in part - the molecular pathogenesis of microcephaly in BS. Finally, our data provide the first evidence of a novel link between BLM dysfunction and transcriptional changes in condensin complex I and II genes. Overall, our study provides novel insights into gene expression profiles in BS on a single cell level linking specific genes and pathways to BLM dysfunction.

## Materials and Methods

### Cell Culture

Two wild-type control fibroblast cell lines (GM08398 and GM00409) and three BS patient fibroblast cell lines (GM02520, GM02932, and GM02548) were purchased from Coriell Institute for Medical Research (see Supplementary Table1 for details). The cells were maintained in Dulbecco’s Modified Eagle Medium (DMEM; Gibco, Life Technologies) supplemented with 10% fetal bovine serum (FBS Superior; Sigma-Aldrich), 100 U/mL penicillin, and 100 μg/mL streptomycin in a 37°C incubator with 5% CO_2_. On the day of single-cell sorting, cells were harvested with 0.05% trypsin-EDTA (Gibco, Life Technologies) and diluted in 1X DPBS before further steps.

### Mutation confirmation of the cell lines

Genomic DNA was extracted from one wild-type control and three patient cell lines using DNeasy Blood and Tissue kit (Qiagen). Specific primers were designed to cover approximately 500 base pairs around the mutations and the regions were PCR-amplified using Qiagen Multiplex PCR kit (Qiagen). The PCR products were sequenced with corresponding primers using canonical Sanger sequencing at SeqLab-Microsynth (Göttingen, Germany) and the obtained results were visualized with the 4Peaks software (66).

### Single-cell transcriptome sequencing

Resuspended cells were stained with Hoechst 33342 and Propidium Iodide (ReadyProbes Cell Viability Imaging Kit, Thermo Fisher) in accordance with the manufacturer’s instructions. After microscope check for proper staining, the cells were counted using CASY cell counter and serial dilutions were prepared to obtain 1 cell/50 nL concentration before single-cell sorting. Single cells were dispensed to a 5184-well microchip using ICELL8 (Takara Bio) system (22) and the well content was evaluated with image processing software of Takara. The wells containing one living single cell were selected with the help of DAPI-positive (one signal spot) and propidium iodide–negative signals. After cell lysis at -80°C, cDNA synthesis was done by SMART-Seq Single Cell kit (Takara Bio) and the library was prepared with Nextera XT DNA library preparation kit (Illumina). The generated library was sequenced on an Illumina HiSeq4000 and the fastq files were generated.

### Quality control, alignment and counting of reads

First, the resulting fastq file was demultiplexed using the demux script of the CogentAP analysis pipeline version 1.0 (Cogent NGS Analysis Pipeline, Takara Bio). The single-end 51 base-pair reads were analyzed by using the analysis part from the CogentAP pipeline. This analysis utilized cutadapt (67) (version 1.18) for trimming of bad quality bases and overrepresented sequences from reads. Afterwards, the reads were aligned against the reference genome GRCh38.12 with Ensembl version 94 using the STAR RNA-Seq aligner (68) (version 2.7.8a). For the counting of gene-level expression values, featureCounts (69) (2.0.1) was used. Cell samples with less than 10,000 reads or less than 300 expressed genes were excluded from the analysis. Additionally, the cells with more counts than three median absolute deviations (MAD) from the total median log10-library size were marked as outliers and were excluded. As the last step, cells with a mitochondrial fraction of 0-0.2 and an intergenic fraction of 0-0.1 were excluded as well.

### Differential gene expression and over-representation analysis

The R-package Limma + Voom (44) for the differential gene expression analysis was used on the filtered and quality-controlled count data. Limma+Voom fits a linear model using the weighted least squares for each gene and combining it with Empirical Bayes smoothing of standard errors. The resulting differentially expressed gene lists were multiple test adjusted (Benjamini-Hochberg) (70) and used as input for the over-representation analysis of Gene-Ontology, KEGG, and MeSH terms using the clusterProfiler (71) package from the R-programming language.

### Western blot

Dermal fibroblast cells were lysed in solution containing 20 mM Tris-HCl (pH 7.5), 150 mM NaCl, 1 mM EDTA (pH 8), 1% NP40, and 2.5% SDS supplemented with protease inhibitors (Halt Protease Inhibitor Cocktail, Thermo Fisher). Protein samples were loaded to a 4-20% Mini-PROTEAN TGX stain-free gel (Bio-Rad), transferred to a PVDF membrane and incubated with respective primary antibodies. The antibodies and the dilutions used in this study were the following: anti-SMC2 (Novus Biologicals, NB100-373), 1:2000; anti-SMC4 (Proteintech, 24758-1-AP), 1:2000; anti-Tubulin (Abcam, ab52866), 1:5000. After incubating the membrane with horseradish peroxidase-conjugated secondary antibodies, the proteins were visualized with enhanced chemiluminescence detection (WesternBright ECL HRP substrate, Advansta) method.

## Acknowledgement

We thank Gabriela Salinas, Orr Shomroni, and Fabian Ludewig from the NGS Integrative Core Unit (Institute of Human Genetics, University Medical Center Göttingen) for generating the raw data of scRNAseq. We are highly grateful to Karin Boss for critically reading the manuscript. This work was supported by the Deutsche Forschungsgemeinschaft (DFG, German Research Foundation) under Research Group FOR 2800 **“**Chromosome Instability: Cross-talk of DNA replication stress and mitotic dysfunction”, SP5 and SPZ to B. Wollnik and Germany’s Excellence Strategy, Cluster of Excellence “Multiscale Bioimaging: from Molecular Machines to Networks of Excitable Cells” (MBExC; EXC 2067/1-390729940) to B. Wollnik.

## Data Availability Statement

The data that support the findings of this study are available on request from the corresponding author. The data are not publicly available due to privacy or ethical restrictions.

## Abbreviations

BS: Bloom syndrome
DSB: double stranded break
sc: single-cell
scRNAseq: single-cell transcriptome sequencing
WT: wild type
QC: quality control
UMAP: Uniform Manifold Approximation and Projection
ORA: over-representation analysis
GO: gene ontology
MeSH: Medical Subject Headings
FA: Fanconi anemia

## Supplementary Figures

**Figure S1.**
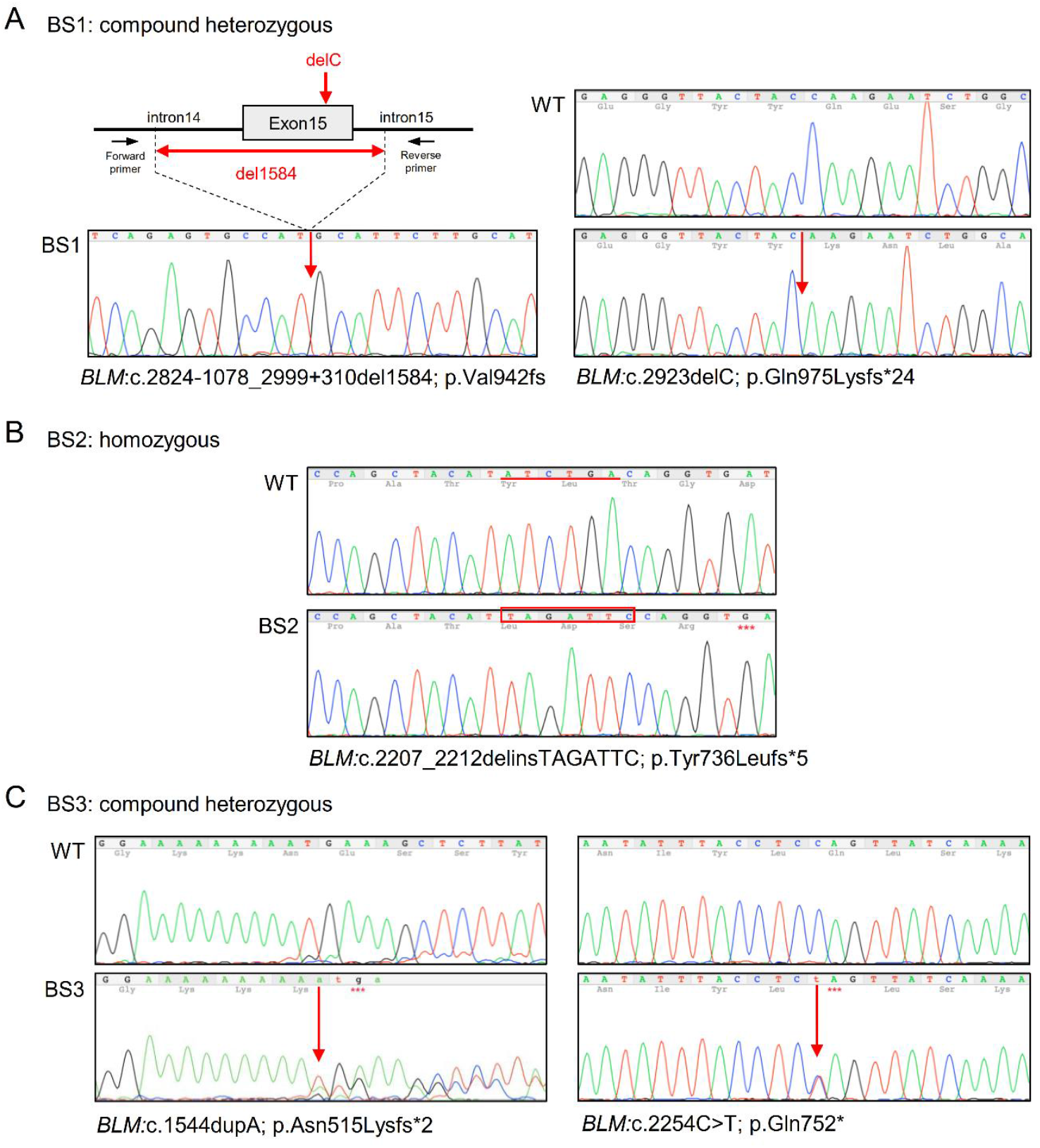
*BLM* gene mutations of the cell lines used in this study were confirmed with Sanger sequencing. (A) BS1 cell line was compound heterozygous, (B) BS2 cell line was homozygous, and (C) BS3 cell line was compound heterozygous for the corresponding loss-of-function mutations in the *BLM* gene. Red arrows indicate the positions of the mutations on the alleles of Bloom syndrome patient cell lines on part (A) and (C). Red underlined sequence shows the deleted sequence on wild-type whereas the sequence marked with red rectangle shows the inserted seqeunce on the BS patient cell line on part (B). Nomenclature is according to *BLM* transcript NM_000057.4.

**Figure S2.**
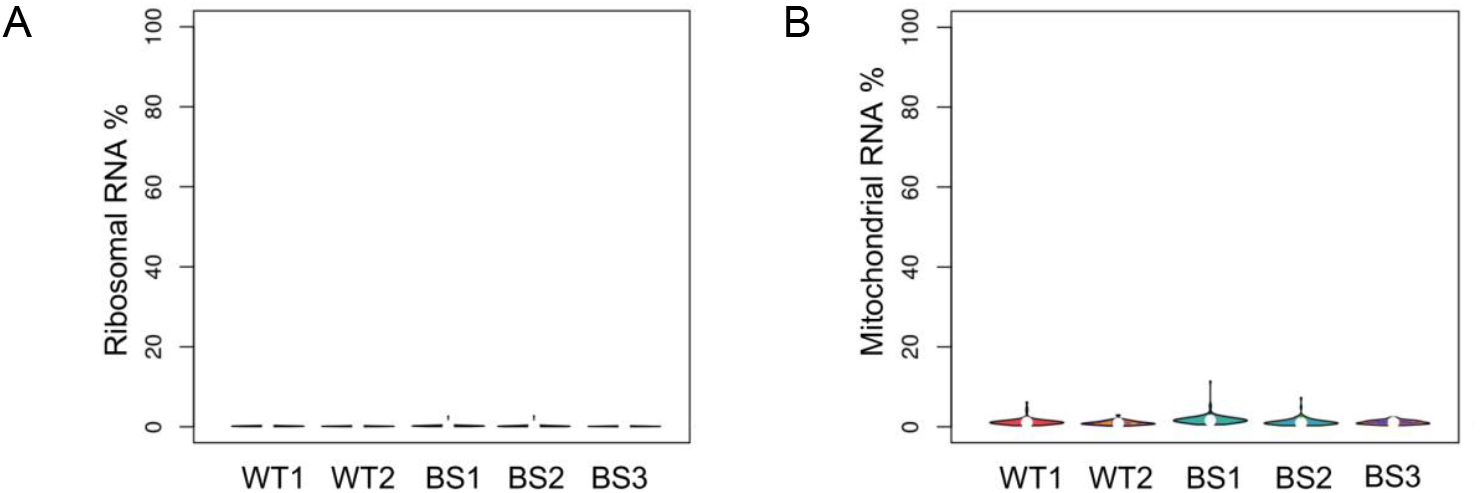
Additional quality control elements support the robustness of the single-cell transcriptome sequencing technique. (A) Ribosomal RNA percentage shows almost no ribosomal RNA detected in the raw data of all samples. (B) Mitochondrial RNA percentage shows less than 10% of overall mitochondrial gene expression among all genes for all samples.

**Figure S3.**
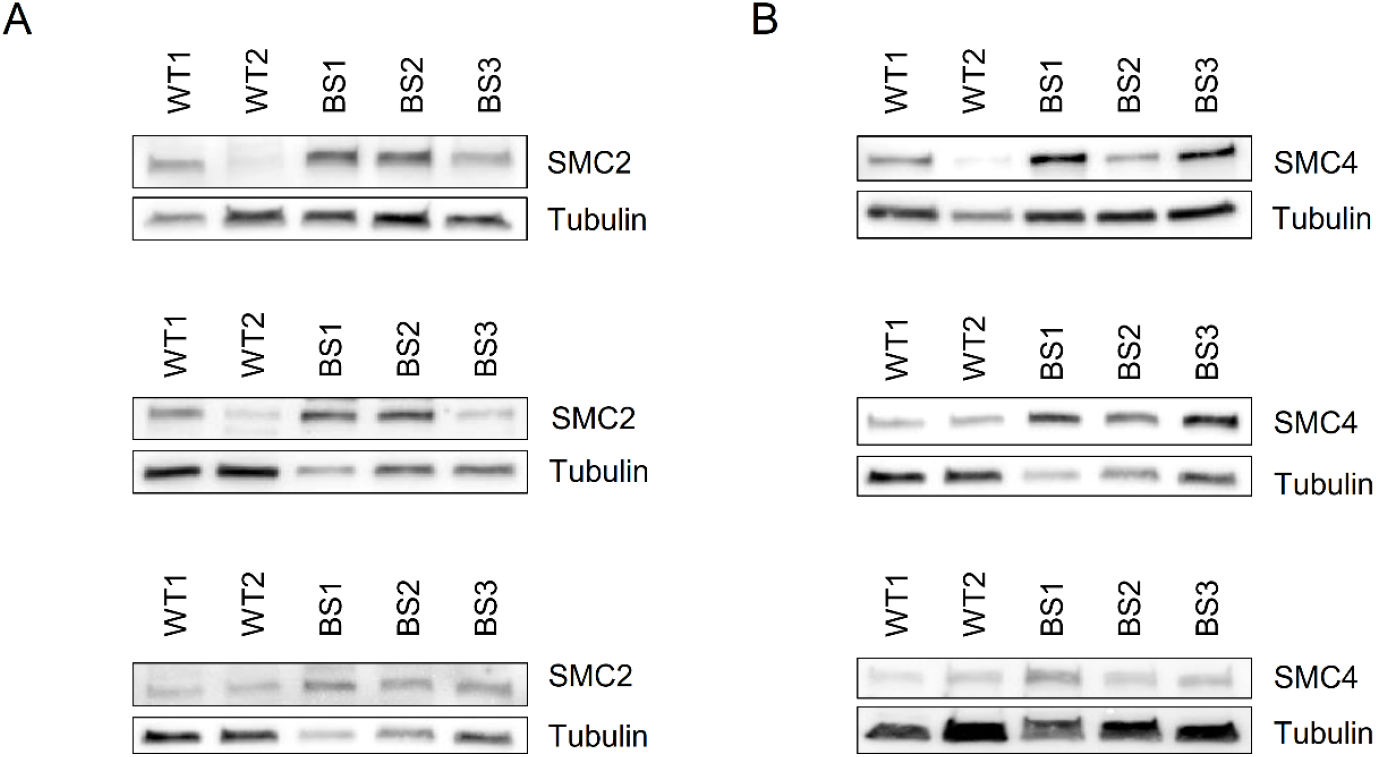
Western blots show the existence of proteins on the cell lysates of fibroblast cell lines used in this study. (A) *SMC2*, (B) *SMC4* genes were expressed on all of the cell lysates. Higher expression levels of SMC2 and SMC4 proteins in the BS cell lines (BS1, BS2, and BS3) can be observed in comparison to control cell lines (WT1 and WT2). The experiments were done in triplicates and the triplicates are given in rows.

## Supplementary Tables

**Supplementary Table 1:** Cell line details used in the study

| Cell line name in the study | Cell line name at Coriell Institute | Age at sampling | Gender | Genotype |
| --- | --- | --- | --- | --- |
| WT1 | GM08398 | 8 | male | wild type |
| WT2 | GM00409 | 7 | male | wild type |
| BS1 | GM02520 | 10 | female | *BLM*:c.2923delC; p.Gln975Lysfs*24 / c.2824-1078_2999+310del1584; p.Val942fs |
| BS2 | GM02932 | 28 | male | homozygous *BLM*:c.2207_2212delinsTAGATTC; p.Tyr736Leufs*5 |
| BS3 | GM02548 | 6 | male | *BLM*:c.1544dupA; p.Asn515Lysfs*2 / c.2254C>T; p.Gln752* |

Nomenclature is according to transcript NM\_000057.4 for *BLM*.

Supplementary Table 2: Differentially expressed genes in BS cells in comparison to WT

Supplementary Table 3: Genes that are included in Figure 3 and the corresponding values

